## Supplementary Materials for "Immersive NREM dreaming preserves subjective sleep depth against declining sleep pressure"

for

**Affiliations**:
[1] *IMT School for Advanced Studies Lucca, MoMiLab Research Unit, Lucca, Italy*
[2] *Department of Biomedical, Metabolic, and Neuronal Sciences, University of Modena and Reggio-Emilia, Modena, Italy*
[3] *University of Padova, Department of General Psychology, Padova, Italy*
[4] *Fondazione Toscana G. Monasterio, Pisa, Italy*
[5] *Sant'Anna School for Advanced Studies, Pisa, Italy*

*** Corresponding author**


† These authors contributed equally to this work

### Supplementary Figure S1

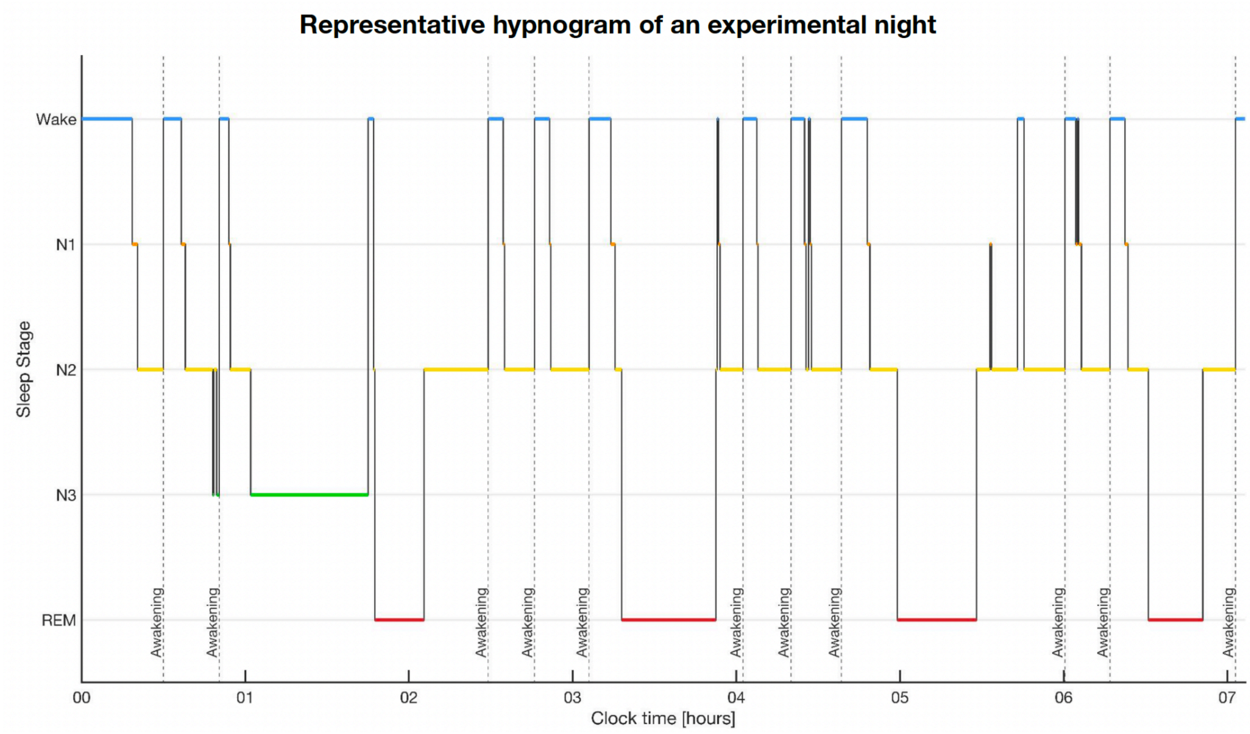

*Figure S1. Representative hypnogram from a single experimental night. The trace illustrates the progression of sleep stages across the night of a participant in Experiment 1, with the x-axis indicating clock time (00 = midnight) and the y-axis denoting sleep stage (Wake, N1, N2, N3, REM). Sleep scoring was performed using U-Sleep. Vertical dashed lines mark experimentally induced awakenings delivered during N2 sleep. By restricting experimental interventions to N2, we maximized data collection within a single stage while minimizing potential alterations to the overall sleep architecture and reducing the risk of cross-stage carryover or interaction effects.*

**Supplementary Figure S2**

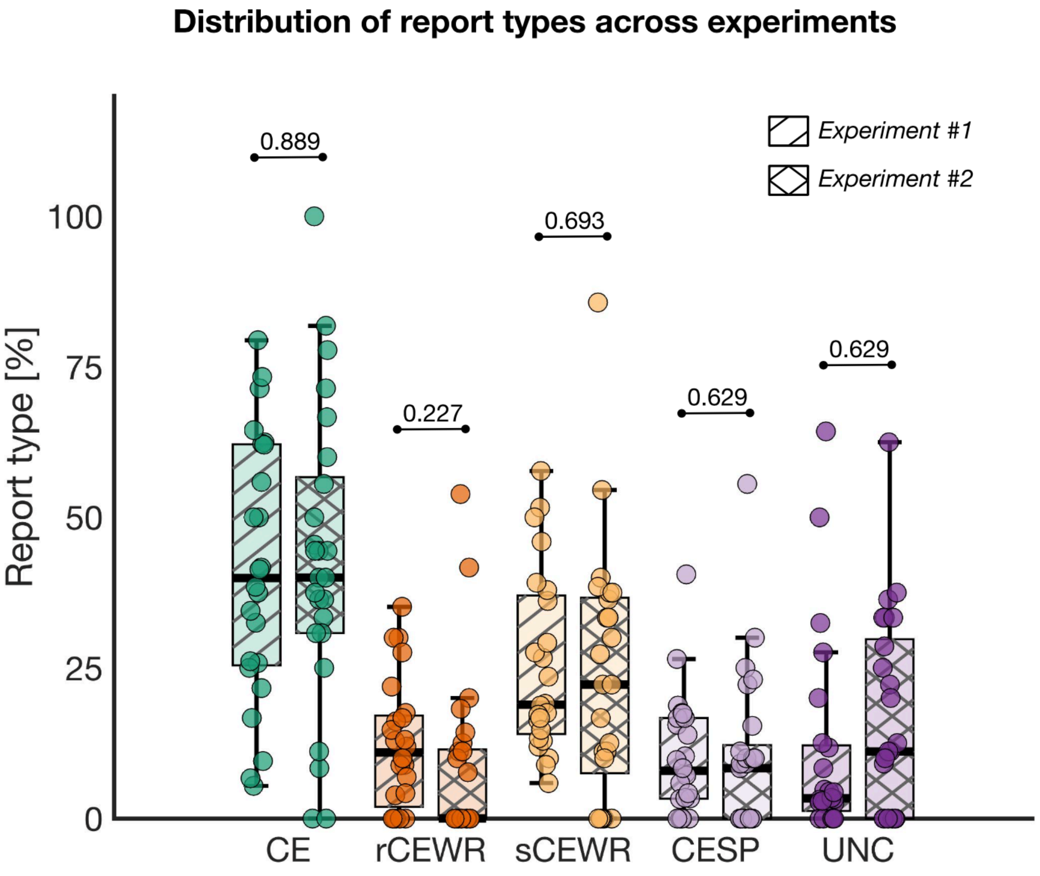

*Figure S2. Distribution of report types across participants and experiments. Boxplots show the percentage of each report type for individual participants in Experiment 1 (single oblique line) and Experiment 2 (crossed lines). For each report type, rank-sum tests were used to assess differences between experiments; FDR-corrected q-values are reported above each pair of boxplots. No significant differences were observed across experiments. In box plots, the box spans the interquartile range (IQR), the horizontal line indicates the median, and whiskers extend to the most extreme values within 1.5 × IQR.. Report type abbreviations: CE, conscious experience; rCEWR, rich conscious experience without recall of content; sCEWR, simple conscious experience without recall of content; CESP, minimal conscious experience with a sense of presence; UNC, unconsciousness.*

**Supplementary Figure S3**

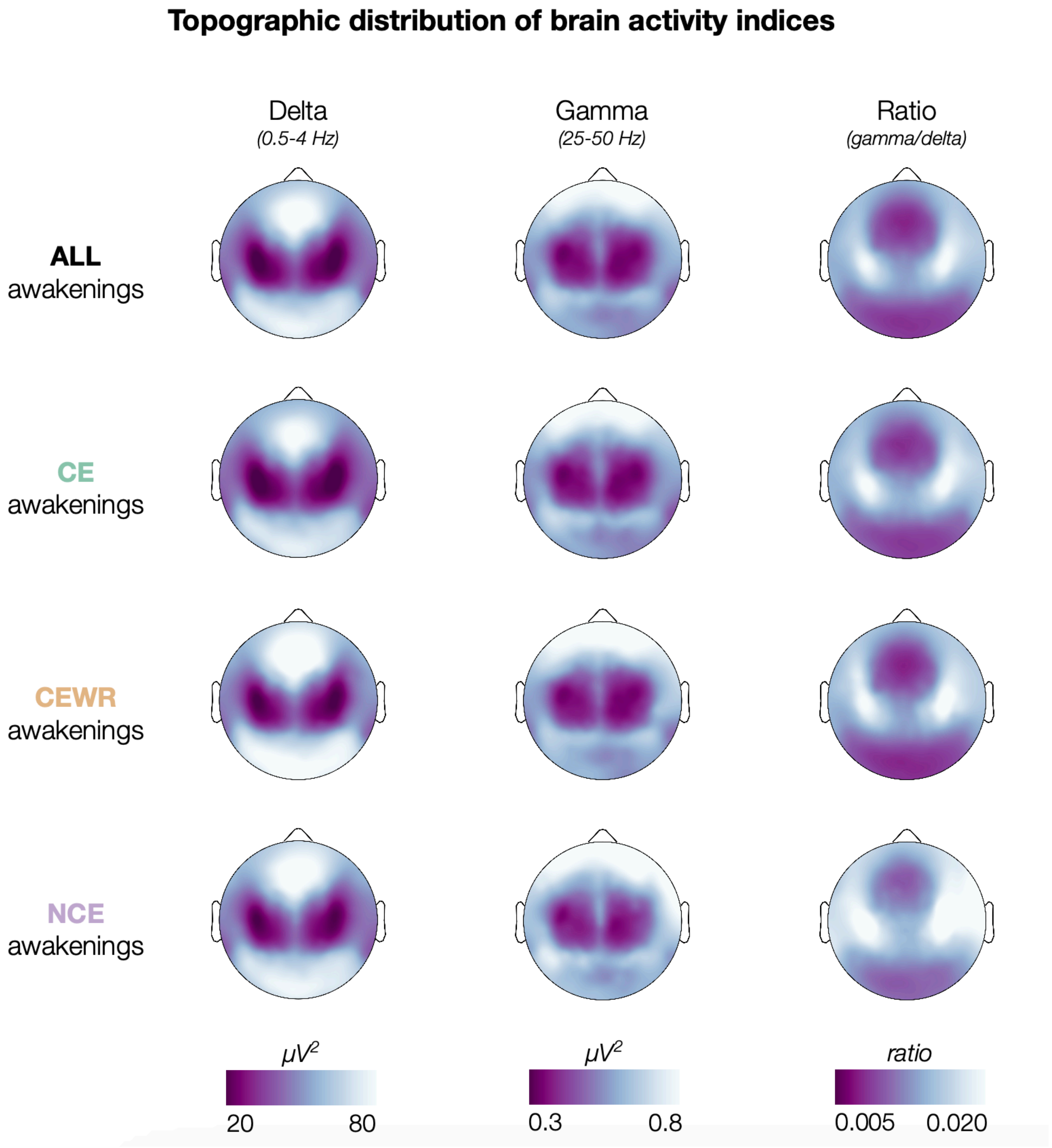

*Figure S3. Topographic distribution of brain activity indices. Topographic maps display the average spatial distribution of EEG indices in the 120 s preceding each awakening, separately for all reports, CE reports, CEWR reports, and NCE reports. Values were first averaged across awakenings within participant, and then across participants, irrespective of experimental condition.*

**Supplementary Figure S4**

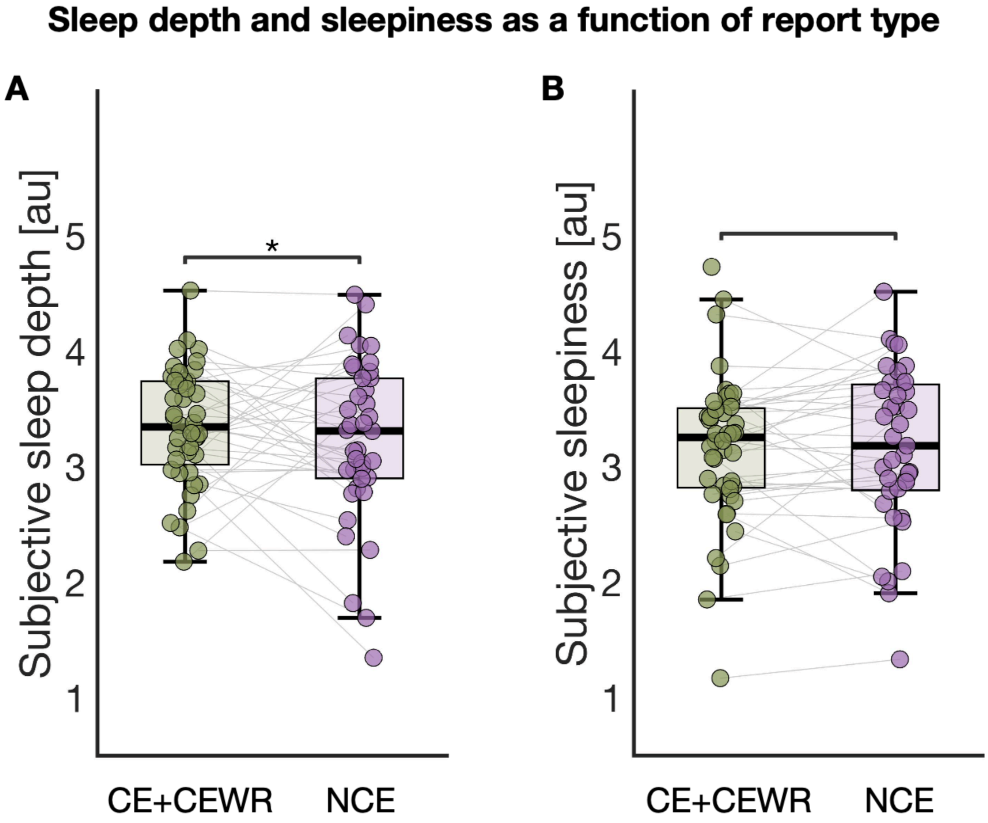

*Figure S4. Relationship between conscious experience and self-reported sleep depth and sleepiness. Participants rated sleep depth (A) and sleepiness (B) on a 5-point Likert scale. Each dot represents the average score for an individual participant. Displayed values are adjusted for experiment, night, and time of night. Asterisks indicate statistical significance based on GLME results: * p < 0.05, ** p < 0.01, *** p < 0.001. In box plots, the box spans the interquartile range (IQR), the horizontal line indicates the median, and whiskers extend to the most extreme values within 1.5 × IQR. Grey lines link data from the same participant across conditions. Color coding: CE+CEWR (reports of conscious experience) = dark green; NCE (no conscious experience) = purple.*

**Supplementary Figure S5**

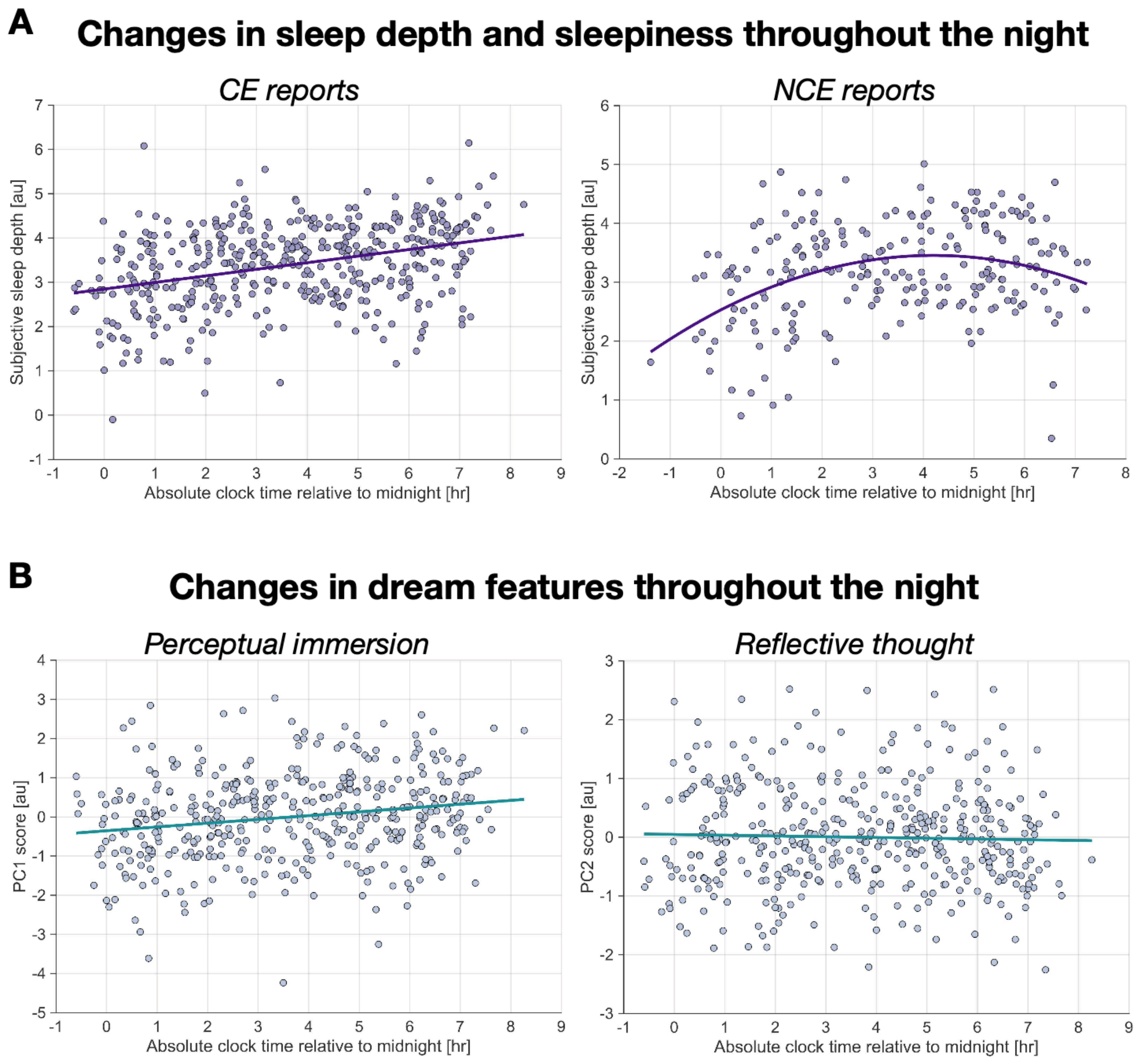

*Figure S5. Time-of-night effects on subjective sleep depth and dream features. (A) Subjective sleep depth as a function of clock time relative to midnight plotted separately for CE reports (left) and NCE reports (right). (B) Dream perceptual immersion (PC1; left) and reflective thought (PC2; right) components plotted as a function of time of night. All values are adjusted for experiment, night, and participant. In panel A, sleep depth is additionally adjusted for sleepiness scores to account for their shared variance. The left plot in panel B is the same shown in Figure 4 of the main text, presented here for comparison with the other plots. Each dot represents one observation (N = 1024 in panel A; N = 432 in panel B, left; N = 228 in panel B, right). Curves represent the best-fitting polynomial model (linear, quadratic, or cubic) selected using the Bayesian Information Criterion (BIC).*

### Supplementary Table S1

| | Experiment #1 | Experiment #2 |
|---|---|---|
| **Participants** [No.] | 24 | 25 |
| **Sex** [No. females/males] | 13/11 | 12/13 |
| **Age** [years] | 26.0 ± 2.3 | 26.6 ± 4.8 |
| **Total Sleep Time** [min] | 330.07 ± 47.14 | 323.82 ± 48.41 |
| **Stage W** [%] | 21.73 ± 10.83 | 22.10 ± 11.00 |
| **Stage N1** [%] | 6.76 ± 4.18 | 6.35 ± 2.99 |
| **Stage N2** [%] | 40.38 ± 8.46 | 44.3 ± 7.56 |
| **Stage N3** [%] | 13.95 ± 5.79 | 12.41 ± 6.1 |
| **Stage REM** [%] | 17.18 ± 6.41 | 14.84 ± 5.71 |
| **Total Awakenings** [No.] | 877 | 260 |
| **Retained Awakenings** [No. (%)] | 783 (89.28%) | 241 (92.69%) |
| **CE reports** [No. (%)] | 330 (42.15%) | 102 (42.32%) |
| **CEWR reports** [No. (%)] | 284 (36.27%) | 80 (33.20%) |
| **NCE reports** [No. (%)] | 169 (21.58%) | 59 (24.48%) |

*Table S1. Characteristics of the samples in the two experiments. The table includes summary statistics regarding the participants' sleep structure (total sleep time and percentages of the different sleep stages) and the performed awakenings (absolute number and percentages of the different report types with respect to the total number of retained awakenings). For the age and the sleep structure indices, values are reported as mean* ± standard deviation.

### Supplementary Table S2

| Model | Predictor | Elec. β | N. Obs. | $R^2$ Adj. | Model p | Coeff. β | CI low | CI high | Coeff. p |
|---|---|---|---|---|---|---|---|---|---|
| All reports | Delta | min | 1024 | - | - | - | - | - | n.s. |
| | | max | 1024 | - | - | - | - | - | n.s. |
| | Gamma | min | 1024 | 0.23144 | < 0.0001 | -0.19225 | -0.31756 | -0.06694 | 0.0026726 |
| | | max | 1024 | 0.25788 | < 0.0001 | -0.51115 | -0.67080 | -0.35150 | 4.9242e-10 |
| | Ratio | min | 1024 | 0.22935 | < 0.0001 | -0.11152 | -0.18828 | -0.03477 | 0.0044421 |
| | | max | 1024 | 0.24734 | < 0.0001 | -0.25416 | -0.34451 | -0.16381 | 4.3007e-08 |
| Interact. | Delta | min | 1024 | 0.23754 | < 0.0001 | -0.30463 | -0.51261 | -0.09664 | 0.0041358 |
| | | max | 1024 | 0.23695 | < 0.0001 | -0.33790 | -0.55185 | -0.12396 | 0.0019936 |
| | Gamma | min | 1024 | 0.24220 | < 0.0001 | 0.38006 | 0.13412 | 0.62600 | 0.0024873 |
| | | max | 1024 | 0.25374 | < 0.0001 | 0.47545 | 0.20508 | 0.74581 | 0.00058191 |
| | Ratio | min | 1024 | 0.25412 | < 0.0001 | 0.23367 | 0.07473 | 0.39260 | 0.0039966 |
| | | max | 1024 | 0.26181 | < 0.0001 | 0.33646 | 0.14986 | 0.52306 | 0.00042106 |
| CE + CEWR reports | Delta | min | 796 | - | - | - | - | - | n.s. |
| | | max | 796 | - | - | - | - | - | n.s. |
| | Gamma | min | 796 | 0.25855 | < 0.0001 | -0.18085 | -0.30602 | -0.05568 | 0.004682 |
| | | max | 796 | 0.27750 | < 0.0001 | -0.43238 | -0.60995 | -0.25481 | 2.092e-06 |
| | Ratio | min | 796 | 0.25966 | < 0.0001 | -0.13032 | -0.21960 | -0.04103 | 0.0042809 |
| | | max | 796 | 0.26738 | < 0.0001 | -0.20357 | -0.30458 | -0.10255 | 8.3201e-05 |
| NCE reports | Delta | min | 228 | - | - | - | - | - | n.s. |
| | | max | 228 | - | - | - | - | - | n.s. |
| | Gamma | min | 228 | 0.23146 | < 0.0001 | -0.35411 | -0.58257 | -0.12565 | 0.0025299 |
| | | max | 228 | 0.25126 | < 0.0001 | -0.71796 | -1.03270 | -0.40320 | 1.1216e-05 |
| | Ratio | min | 228 | 0.22979 | < 0.0001 | -0.23383 | -0.39447 | -0.07318 | 0.0045244 |
| | | max | 228 | 0.26163 | < 0.0001 | -0.41702 | -0.60060 | -0.23344 | 1.214e-05 |

*Table S2. Results of the GLME analyses examining the relationship between brain activity and subjective sleep depth at the electrode level. The first two sets of models tested: (1) the main effects of neural predictors (delta power, gamma power, gamma/delta ratio), and (2) interactions between each neural predictor and report type (CE+CEWR vs. NCE). The third and fourth sets of models examined the main effects of neural predictors separately for CE+CEWR (3) and NCE (4) reports. For each analysis yielding significant effects, the table presents results from the electrodes with the smallest and largest absolute β coefficients among those reaching significance. All models included experiment, night, and time of night as fixed effects, and participant as a random effect. Reported metrics include the number of observations (N. Obs.), adjusted model $R^2$ ($R^2$ Adj.), model p-value relative to a null model with only the random effect (Model p), the estimated effect size (Coeff. β), its 95% confidence interval (Coeff. CI low and high), and associated p-value (Coeff. p).*

### Supplementary Table S3

| Model | Predictor | Elec. β | N. Obs. | $R^2$ Adj. | Model p | Coeff. β | CI low | CI high | Coeff. p |
|---|---|---|---|---|---|---|---|---|---|
| **All reports** | Delta | min | 1024 | - | - | - | - | - | n.s. |
| | | max | 1024 | - | - | - | - | - | n.s. |
| | Gamma | min | 1024 | 0.44365 | < 0.0001 | -0.15113 | -0.24792 | -0.05435 | 0.0022405 |
| | | max | 1024 | 0.45216 | < 0.0001 | -0.31475 | -0.44850 | -0.18101 | 4.3682e-06 |
| | Ratio | min | 1024 | 0.44351 | < 0.0001 | -0.09082 | -0.15248 | -0.02917 | 0.0039262 |
| | | max | 1024 | 0.44598 | < 0.0001 | -0.13414 | -0.20571 | -0.06256 | 0.00024786 |
| **Interact.** | Delta | min | 1024 | - | - | - | - | - | n.s. |
| | | max | 1024 | - | - | - | - | - | n.s. |
| | Gamma | min | 1024 | - | - | - | - | - | n.s. |
| | | max | 1024 | - | - | - | - | - | n.s. |
| | Ratio | min | 1024 | - | - | - | - | - | n.s. |
| | | max | 1024 | - | - | - | - | - | n.s. |

*Table S3. Results of the GLME analyses examining the relationship between brain activity and subjective sleepiness at the electrode level. The first two sets of models tested: (1) the main effects of neural predictors (delta power, gamma power, gamma/delta ratio), and (2) interactions between each neural predictor and report type (CE+CEWR vs. NCE). Since no significant interactions were found, we did not examine the main effects of neural predictors separately for CE+CEWR and NCE reports. For each analysis yielding significant effects, the table presents results from the electrodes with the smallest and largest absolute β coefficients among those reaching significance. All models included experiment, night, and time of night as fixed effects, and participant as a random effect. Reported metrics include the number of observations (N. Obs.), adjusted model $R^2$ ($R^2$ Adj.), model p-value relative to a null model with only the random effect (Model p), the estimated effect size (Coeff. β), its 95% confidence interval (Coeff. CI low and high), and associated p-value (Coeff. p).*

### Supplementary Table S4

| Contrast | N. Obs. | $R^2$ Adj. | Model p | Coeff. β | CI low | CI high | Coeff. p | Coeff. q |
|---|---|---|---|---|---|---|---|---|
| (CE+CEWR) vs. NCE | 1024 | 0.2281 | < 0.0001 | -0.18317 | -0.32698 | -0.039353 | **0.012602** | - |
| CE vs. CEWR | 796 | 0.25136 | < 0.0001 | 0.01223 | -0.12036 | 0.14483 | 0.85636 | 2.0981 |
| CE vs. CESP | 538 | 0.21447 | < 0.0001 | -0.55372 | -0.77180 | -0.33564 | **8.2845e-07** | **< 0.0001** |
| CE vs. UNC | 554 | 0.19783 | < 0.0001 | 0.18280 | -0.03964 | 0.40524 | 0.10705 | 0.3147 |
| CEWR vs. CESP | 470 | 0.33161 | < 0.0001 | 0.55475 | 0.36616 | 0.74333 | **1.3693e-08** | **< 0.0001** |
| CEWR vs. UNC | 486 | 0.26573 | < 0.0001 | 0.16908 | -0.02950 | 0.36765 | 0.094966 | 0.3147 |
| CESP vs. UNC | 228 | 0.2662 | < 0.0001 | 0.59411 | 0.32292 | 0.86530 | **2.3831e-05** | **0.0001** |
| rCEWR vs. sCEWR | 361 | 0.31495 | < 0.0001 | -0.25961 | -0.46033 | -0.05889 | **0.011393** | - |

*Table S4. Results of GLME analyses testing whether subjective sleep depth differed across conscious states during sleep. Each model compares a specific pair of report types (see 'Contrast' column) and includes experiment, night, and time of night as fixed effects, and participant as a random effect. Reported metrics include the number of observations (N. Obs.), adjusted $R^2$ ($R^2$ Adj.), model p-value compared to a null model with only the random effect (Model p), estimated effect size (Coeff. β), its 95% confidence interval (Coeff. CI low and high), and p-value for the effect (Coeff. p). For comparisons involving the four-level classification (CE, CEWR, CESP, UNC; Figure 2), false discovery rate (FDR) correction was applied, and the resulting q-value is shown in the final column. Statistically significant effects ($q < 0.05$) are indicated in bold.*

### Supplementary Table S5

| Contrast | N. Obs. | $R^2$ Adj. | Model p | Coeff. β | CI low | CI high | Coeff. p | Coeff. q |
|---|---|---|---|---|---|---|---|---|
| (CE+CEWR) vs. NCE | 1024 | 0.43855 | < 0.0001 | 0.06651 | -0.049595 | 0.18261 | 0.26124 | - |
| CE vs. CEWR | 796 | 0.44764 | < 0.0001 | 0.058982 | -0.048759 | 0.16672 | 0.28287 | 0.6930 |
| CE vs. CESP | 538 | 0.30247 | < 0.0001 | -0.17803 | -0.35543 | -0.00063 | 0.049198 | 0.1446 |
| CE vs. UNC | 554 | 0.31639 | < 0.0001 | 0.28859 | 0.10541 | 0.47177 | **0.0020711** | **0.0152** |
| CEWR vs. CESP | 470 | 0.58057 | < 0.0001 | 0.19431 | 0.035204 | 0.35341 | 0.016793 | 0.0617 |
| CEWR vs. UNC | 486 | 0.5927 | < 0.0001 | 0.31057 | 0.14732 | 0.47381 | **0.0002077** | **0.0031** |
| CESP vs. UNC | 228 | 0.46278 | 0.0005 | 0.33026 | 0.089618 | 0.57091 | **0.0073694** | **0.0361** |
| rCEWR vs. sCEWR | 361 | 0.61957 | < 0.0001 | -0.095761 | -0.26543 | 0.073912 | 0.26777 | - |

*Table S5. Results of GLME analyses testing whether subjective sleepiness differed across conscious states during sleep. Each model compares a specific pair of report types (see 'Contrast' column) and includes experiment, night, and time of night as fixed effects, and participant as a random effect. Reported metrics include the number of observations (N. Obs.), adjusted $R^2$ ($R^2$ Adj.), model p-value compared to a null model with only the random effect (Model p), estimated effect size (Coeff. β), its 95% confidence interval (Coeff. CI low and high), and p-value for the effect (Coeff. p). For comparisons involving the four-level classification (CE, CEWR, CESP, UNC; Figure 2), false discovery rate (FDR) correction was applied, and the resulting q-value is shown in the final column. Statistically significant effects ($q < 0.05$) are indicated in bold.*

### Supplementary Table S6

| Predictor | Elec. β | N. Obs. | $R^2$ Adj. | Model p | Coeff. β | CI low | CI high | Coeff. p |
|---|---|---|---|---|---|---|---|---|
| **Sleep depth** | PC1 | 427 | 0.29968 | < 0.0001 | 0.27689 | 0.20777 | 0.34601 | **2.9597e-14** |
| | PC2 | 427 | 0.24391 | < 0.0001 | -0.19473 | -0.28696 | -0.10249 | **4.0294e-05** |
| **Sleepiness** | PC1 | 427 | 0.25189 | 0.00303 | 0.078884 | 0.017956 | 0.13981 | **0.011288** |
| | PC2 | 427 | 0.24241 | 0.02925 | -0.03227 | -0.10882 | 0.044277 | 0.40777 |

*Table S6. Results of the GLME analyses examining the relationship between the phenomenological features of conscious experience and subjective sleep depth or sleepiness. All models included experiment, night, and time of night as fixed effects, and participant as a random effect. Reported metrics include the number of observations (N. Obs.), adjusted model $R^2$ ($R^2$ Adj.), model p-value relative to a null model with only the random effect (Model p), the estimated effect size (Coeff. β), its 95% confidence interval (Coeff. CI low and high), and associated p-value (Coeff. p). Statistically significant effects ($p < 0.05$) are indicated in bold.*

### Supplementary Table S7

| **Predicted var.** | **N. Obs.** | **$R^2$ Adj.** | **Model p** | **Coeff. β** | **CI low** | **CI high** | **Coeff. p** |
|---|---|---|---|---|---|---|---|
| Seep depth | 1024 | 0.35496 | < 0.0001 | 0.12713 | 0.1029 | 0.15137 | **1.0412e-23** |
| Sleepiness | 1024 | 0.53312 | < 0.0001 | -0.012044 | -0.032306 | 0.0082172 | 0.2437 |
| Delta power | 1024 | 0.22599 | < 0.0001 | -0.071342 | -0.090281 | -0.052404 | **3.0091e-13** |
| Gamma power | 1024 | 0.40913 | 0.0009 | -0.0068527 | -0.018638 | 0.0049321 | 0.25412 |
| Gamma/Delta ratio | 1024 | 0.27374 | < 0.0001 | 0.064743 | 0.043795 | 0.085692 | **1.8624e-09** |
| PC1 | 427 | 0.30589 | 0.0015 | 0.10339 | 0.05309 | 0.15369 | **6.3486e-05** |
| PC2 | 427 | 0.29589 | 0.7862 | -0.013566 | -0.053961 | 0.02683 | 0.50956 |

*Table S7. Results of GLME analyses investigating the effect of time of the night on sleep depth, sleepiness, brain activity, and dream features. All models include experiment, night, and time of night as fixed effects, and participant as a random effect. To account for shared variance between subjective sleep depth and sleepiness, each of the models exploring these variables included the other measure as an additional fixed effect. Reported metrics include the number of observations (N. Obs.), adjusted $R^2$ ($R^2$ Adj.), model p-value compared to a null model with only the random effect (Model p), estimated effect size (Coeff. β), its 95% confidence interval (Coeff. CI low and high), and p-value for the effect (Coeff. p). Statistically significant effects ($p < 0.05$) are indicated in bold.*

### Supplementary Table S8

| Prompts | Instructions |
|---|---|
| *1. What was on your mind just before waking up?* | We want to know whether you remember having any subjective experience in the moments before the alarm sounded. It is important to note that we are not only referring to classic "dreams," which are typically visual and associated with a story, but to any type of experience, including thoughts, images, sensations, or emotions. If you had no experience, simply report this. Likewise, let us know if you think you had an experience but cannot remember what it was. Depending on your response, you will be asked a follow-up question. |
| *2a. Recall the experience.* | If you remember having an experience, please describe it. Focus on the last experience you had in the moments before the alarm sounded. Tell us what you remember. It is okay if you cannot recall all the details or impressions.<br><br>*Example 1*: You were at a party with friends. Suddenly, wolves arrived, and you ran away. You then found yourself in a forest, stopped to look at a plant with very colorful and fragrant flowers, and thought they were beautiful. At that moment, the alarm sounded. The last experience in this case is related to the flowers and includes visual elements (seeing the flowers and their colors), olfactory elements (smelling the flowers), and cognitive elements (thinking that they were beautiful).<br><br>*Example 2*: You were thinking about your to-do list for the next day when you suddenly remembered an urgent deadline, which made you feel anxious. At that moment, the alarm sounded. The last experience in this case includes both the thought about the deadline and the emotional state of anxiety.<br><br>After describing your last experience, you will be asked to estimate how long you had continuous experiences before the alarm sounded. Again, we refer to experiences of any kind, not necessarily forming a single coherent narrative. You do not need to recall specific details of the experience. You should report, approximately, how long you believe you have had continuous experiences before the alarm sounded, not necessarily in one coherent narrative. It is important to note that this question does not refer to the time that passed since the previous awakening but to how much time was occupied by experiences of some kind. For example, if 25 minutes passed since your last awakening but you believe you had subjective experiences only during the last 10 minutes before the current awakening, your response should be "10 minutes." |
| *2b. Do you have the impression that the experience was rich in details or events?* | If you had an experience but cannot remember it, you will be asked whether you think it was rich in details or events. Sometimes, even if we cannot recall an experience, we may have the impression that it was vivid, long, or complex. In other cases, we may only have a vague sense of having had an experience without any specific impression of its content. Simply answer "yes" (it was rich in details or events) or "no" (it was not). |
| *2c. Did you have the feeling of being "present"?* | If you did not have any experience, you will be asked whether you had a feeling of "presence" before the alarm sounded. By this, we mean a sense of being alive and/or perceiving the present moment and the passage of time. Answer "yes" (I was present) or "no" (I was not). |
| *3. How deeply asleep did you feel before the alarm sounded, on a scale from 1 to 5, where 1 is completely awake and 5 is deeply asleep?* | This question refers to your subjective state just before the alarm sounded. Use only whole numbers in your response. "1" means you felt fully alert and awake, "3" indicates an intermediate state between wakefulness and sleep, and "5" means you felt deeply asleep. |
| *4. How sleepy do you feel, on a scale from 1 to 5, where 1 is not sleepy at all and 5 is extremely sleepy?* | This question refers to your subjective state after the alarm sounded. Use only whole numbers in your response. "1" means you do not feel sleepy or drowsy at all, "3" indicates moderate sleepiness and drowsiness, and "5" means you feel extremely sleepy and struggle to stay awake. |

| | |
|---|---|
| *5. Do you think you perceived a stimulus just before the alarm sounded?* | During sleep, it is sometimes possible to perceive (or believe you perceived) events occurring around you. These could be sensory stimuli presented as part of the experiment, unintentional noises from the researchers, or external factors unrelated to the experiment.<br>In some cases, stimuli might be incorporated into the dream experience while still being recognizable as coming from the external environment (e.g., hearing water running from a faucet might be associated with seeing a waterfall in a dream). Please indicate whether you think you consciously perceived any external stimulus just before the alarm sounded. Do not include stimuli that you believe were generated solely by your own mind. If you perceived something, specify whether it was auditory, visual, tactile, or of another kind (e.g., smell, warmth, pain, etc.). |
| *The following questions refer to experiences (dreams, thoughts, images, sensations, emotions) that occurred just before waking up. They do not refer to any previous experiences, even if these were continuous with the ones happening at the moment of the alarm sound.* | |
| *6. How vivid was the experience on a scale from 1 to 5, where 1 is not vivid at all and 5 is extremely vivid?* | This question refers to the "vividness" of the experience, meaning its level of clarity and sharpness. It is important to note that this question does not only refer to the visual aspect of the experience but to its overall vividness, regardless of its nature, including different types of sensory experiences, thoughts, emotions, or sensations. For example, a very clear and well-defined thought, similar to a waking thought, can be considered a vivid experience. Only whole numbers should be used in the response. A response of "1" means that the experience was unclear and blurry (e.g., a vague and indistinct image), "3" indicates an intermediate level of vividness, while "5" means the experience was extremely sharp and clear (e.g., an image as well-defined as in normal vision). |
| *7. To what extent was the experience perceptual rather than thought-based, on a scale from 1 to 5, where 1 is purely thought and 5 is purely perception?* | This question refers to the type of content in the experience. Our experiences can take the form of abstract thoughts (e.g., thinking about things to do the next day) or sensory-perceptual elements (e.g., seeing a landscape, hearing a voice, perceiving a smell, etc.). Some experiences contain both elements (e.g., seeing an object that reminds you of something you need to do the next day). Only whole numbers should be used in the response. A response of "1" means the experience consisted of pure abstract thought without any sensory-perceptual aspects, "3" indicates that thought and perceptual elements were present in equal measure, while "5" means the experience was purely sensory and perceptual, with no thought or reflection. |
| *8. Did the experience contain sensory content? Visual? Auditory? Tactile? Olfactory? Gustatory?* | Here, you are asked to specify whether the experience contained at least one recognizable sensory element of a visual, auditory, tactile, olfactory, or gustatory nature. You should respond "yes" or "no" to each type of sensory content mentioned by the experimenter. Keep in mind that it is entirely acceptable to respond "no" to all questions, as in the case of experiences consisting purely of thought, without any sensory or perceptual aspects. |
| *9. To what extent was the experience related to real-life elements, situations, or events encountered during wakefulness, on a scale from 1 to 5, where 1 is not related at all and 5 is fully related?* | This question aims to assess how much the experience drew from or referenced memories of elements (e.g., objects, people, places), situations, or events actually encountered during wakefulness. Conscious experiences during sleep can sometimes appear as more or less accurate reworkings of recent (e.g., dreaming about a work meeting that took place that same day or a few days prior) or remote memories (e.g., dreaming about being in a place visited in childhood). Additionally, a single experience may include elements from multiple, seemingly unrelated memories. In other cases, the experience may not reference any specific memory (e.g., dreaming of flying through the clouds). Only whole numbers should be used in the response. A response of "1" means the experience was not directly related to real memories, "3" indicates it was moderately related, while "5" means the experience faithfully reproduced real memories. |
| *10. How bizarre was the experience on a scale from 1 to 5, where 1 is not bizarre at all and 5 is extremely bizarre?* | This question refers to the level of "strangeness" in the experience. Experiences can sometimes include elements that seem odd or "unnatural" (e.g., dreaming of flying on the back of an eagle or talking to someone familiar who appears different than usual). It is important to remember that this question asks you to assess how strange or bizarre the experience felt at the moment of waking up, after having lived it. Only whole numbers should be used in the response. A response of "1" means the experience contained no bizarre elements, "3" indicates an intermediate level of bizarreness, while "5" means you found the experience extremely bizarre. |

| | |
|---|---|
| *11. To what extent were you aware that you were dreaming, on a scale from 1 to 5, where 1 is not aware at all and 5 is fully aware?* | This question refers to your level of awareness that the experience was not real (i.e., it was like a dream). It is important to note that awareness of having a conscious experience during sleep can fluctuate within the experience itself. Sometimes, we temporarily become aware but then forget about it, while other times, we realize it right before waking up. Your response should reflect your level of awareness during the last experience before waking up. Only whole numbers should be used in the response. A response of "1" means there was no awareness of being asleep, "3" indicates a state of uncertainty (e.g., realizing that some aspects of the experience were particularly bizarre but not concluding with certainty that they were unreal), while "5" means there was full awareness. |
| *12. To what extent did you have voluntary control over the content and progression of the experience, on a scale from 1 to 5, where 1 is no control and 5 is full control?* | This question refers to your level of voluntary control over the content and flow of the experience. It is important to note that the level of control can vary within the experience itself. Your response should reflect the degree of control you had during the last experience before waking up. Only whole numbers should be used in the response. A response of "1" means there was no ability to voluntarily control the content or direction of the experience, "3" indicates an intermediate level of control (e.g., the ability to make voluntary choices within the experience but not to alter aspects such as characters, settings, etc.), while "5" means there was full control over the experience, including the ability to modify every aspect. |
| *13. How would you rate the emotional valence of the experience on a scale from 1 to 5, where 1 is very negative and 5 is very positive?* | This question refers specifically to the valence or "emotional tone" of the experience. Our experiences can be predominantly associated with negative, neutral, or positive emotions. Only whole numbers should be used in the response. A response of "1" means the experience was associated with extremely negative emotions (e.g., a bad dream or nightmare), "3" indicates a neutral emotional content (neither predominantly positive nor predominantly negative), while "5" means the experience was associated with extremely positive emotions. It is important to note that the score does not reflect the strength or intensity of emotions but only the subjective valence as more or less negative/positive, or neutral. |
| *14. How would you rate the emotional intensity of the experience on a scale from 1 to 5, where 1 is very weak and 5 is very strong?* | This question specifically refers to the intensity of emotions in the experience. Regardless of their valence, our experiences can be associated with emotional states of varying strength and intensity. Only whole numbers should be used in the response. A response of "1" means the experience was associated with minimal emotional intensity, "3" indicates an intermediate intensity, while "5" means the experience was associated with extremely strong and intense emotions. It is important to remember that the score does not reflect the valence or tone of emotions but only their strength. |
| *15. How confident are you that your memories of the experience are complete and accurate on a scale from 1 to 5, where 1 is not confident at all and 5 is very confident?* | This question refers to your level of certainty that what you remember about the experience accurately and completely reflects what actually occurred. Only whole numbers should be used in the response. A response of "1" means you do not feel confident that you remember the experience completely and accurately, "3" indicates an intermediate level of confidence, while "5" means you feel certain that you remember the experience completely and accurately. In other words, a lower score should be given if you believe you have forgotten parts of the experience or are unsure about remembering certain details correctly. |

***Table S13.*** *Dream questionnaire. Instructions for the volunteers, translated from Italian.*

### Supplementary Text - Additional experimental procedures

#### MRI data acquisition

All participants underwent an MRI session that included the acquisition of high-resolution anatomical images using a magnetization-prepared rapid gradient echo (MPRAGE) sequence (TR = 7 ms, TE = 3.2 ms, flip angle = 9°, field of view = 180 mm, acquisition matrix = 224 × 224, voxel size = 1 × 1 × 1 $mm^3$, 180 sagittal slices). Scans were acquired on a Philips 3T Ingenia system equipped with a 32-channel phased-array head coil. MRI data were not included in the present report.

#### Experiment #1: sensory stimulation before sleep

Participants arrived at the sleep laboratory around 6.30 PM. After EEG cap preparation, they completed a two-hour task session involving either a visual, auditory, or tactile stimulus duration discrimination task or a battery of standardized activities involving all three sensory modalities. Each experimental night was preceded by only one of these tasks, and their order was randomized across participants. Before and after the task session, participants completed three two-minute-long resting state hd-EEG recordings with their eyes open.

In the duration discrimination task, participants were presented with pairs of tones, vibrotactile pulses, or light flashes and asked to judge whether they had identical or different duration. While seated in front of a computer screen, participants maintained fixation on a central black cross throughout the task. To maintain participants' engagement, the duration difference between stimulus pairs was progressively reduced across blocks to increase the difficulty level, with minimum differences set at 400 ms, 200 ms, 100 ms, or 50 ms. Individual stimulus durations ranged from 200 to 800 ms in 50 ms increments, and each pair of stimuli was separated by a 500 ms inter-stimulus interval. Participants responded using designated keys on a numeric keypad. After each given response, a new pair of stimuli was presented. Each difficulty level included two task blocks with 30 stimuli (20% with matched duration). Participants were instructed to respond as fast and as accurately as possible. Feedback on their performance was provided after each stimulus pair by changing the color of the fixation cross to green (correct) or red (wrong) for 500 ms. An additional feedback indicating the percentage of correct responses was shown on the computer screen at the end of each block. Visual stimuli consisted of flashes of the monitor's background obtained by changing the color from gray to white. Auditory stimuli were 1000 Hz tones including five-ms fade-in and fade-out ramps presented through in-hear headphones (*Maxrock*, Guangdong, China). Mechanical vibratory stimuli (80 Hz) were presented using a dedicated stimulator placed on the index finger of the dominant hand (*Tactamp* and *Tactors*, *Dancer Design*, Ingleton, UK).

The standardized activities were selected to engage auditory, visual, and tactile sensory modalities. In the "*guess the sound*" challenge, participants were presented with 50 different sounds and had to guess their source. Similarly, in the "*guess the soundtrack*" challenge they had to recognize 42 famous movies and series from their soundtrack. Other activities included solving a set of wooden puzzles, building a LEGO dinosaur, completing a "*word search*" game, and completing a large "*connect the dots*" game. The order of the activities was randomized across participants.

**Experiment #2: sensory stimulation during sleep**

Participants arrived at the sleep laboratory around 8.30 PM. After the EEG cap and the peripheral sensors were mounted, three different stimulation devices were placed on each participant. Specifically, the same vibrotactile stimulator used in Experiment #1 was taped on the index finger of the dominant hand. Moreover, participants were asked to wear in-ear headphones and a modified sleep mask including two red LED lights (*Mallory Sonalert Products Inc.*, IN, US). The headphones were taped to the EEG cap to avoid the risk of them sliding out during the night. The LEDs were positioned bilaterally in 8 mm cutouts in the sleep mask centered above the eyes. They emitted light with a wavelength of 622 nm and an intensity level of 1500 millicandela. These devices were used to deliver single-pulse 50-ms stimulations during the night. Auditory stimuli consisted of 1000 Hz pure tones (including a five-ms ramp-up and five-ms ramp-down) at a stable intensity level of 40 dB. Tactile stimuli were 80 Hz mechanical vibrations, while visual stimuli consisted of a single red light flash. These stimuli were presented in pseudo-random order during N2 sleep with the aim of inducing a K-complex. Awakenings were induced 4 to 6 seconds after stimuli that successfully evoked a K-complex. In addition to stimulation trials, we also performed sham awakenings not preceded by a stimulation. Of note, if a stimulus failed to evoke a K-complex, the stimulation was repeated once more after a minimum of 2 minutes. If a KC was not evoked by the repeated stimulus, a different stimulation modality or a sham awakening were performed after at least 2 minutes to avoid habituation. Thus, sham awakenings were never closely preceded by stimulation. Only sham trials were analyzed in the present study.

Prior to the beginning of the sleep recording, resting-state recordings with eyes closed were acquired in three two-minute-long blocks.
